# Identification of candidate genes controlling red seed coat color in cowpea (*Vigna unguiculata* [L.] Walp)

**DOI:** 10.1101/2024.01.08.574663

**Authors:** Ira A. Herniter, María Muñoz-Amatriaín, Sassoum Lo, Yi-Ning Guo, Stefano Lonardi, Timothy J. Close

## Abstract

Seed coat color is an important consumer-related train in cowpea (*Vigna unguiculata* [L.] Walp.) and has been a subject of study for over a century. Utilizing newly available resources, including mapping populations, a high-density genotyping platform, and several genome assemblies, red seed coat color has been mapped to two loci, *Red-1* (*R-1*) and *Red-2* (*R-2*), on Vu03 and Vu07, respectively. A gene model (*Vigun03g118700*) encoding a dihydroflavonol 4-reductase, a homolog of anthocyanidin reductase 1, which catalyzes the biosynthesis of epicatechin from cyanidin, has been identified as a candidate gene for *R-1*. Possible causative variants have been also identified for *Vigun03g118700*. A gene model on Vu07 (*Vigun07g118500*), with predicted nucleolar function and high relative expression in the developing seed, has been identified as a candidate for *R-2*. The observed red color is believed to be the result of a buildup of cyanidins in the seed coat.

## 1. Introduction

Cowpea (*Vigna unguiculata* [L.] Walp.) is a diploid (2n = 22) warm season legume which is primarily grown in sub-Saharan Africa, where it serves as a major source of protein and calories. Further production occurs in the Mediterranean Basin, southeast Asia, Latin America, and the United States. Just over 8.9 million metric tonnes of dry cowpeas were reported worldwide in 2019 [1], though these numbers do not include Brazil, Australia, and some other relatively large producers. Most of the production in sub-Saharan Africa is by smallholder farmers in marginal conditions, often as an intercrop with maize, sorghum, or millet [2]. Due to its high adaptability to both heat and drought and its association with nitrogen fixing bacteria, cowpea is a versatile crop [2], [3].

The most common form of consumption of cowpea is as a dry grain, with seeds used whole or ground into flour [4], [5]. Seed coat color is an important consumer-related trait in cowpea, as consumers usually make decisions about the quality and presumed taste of a product based on appearance [6], [7]. Cowpea grain displays a variety of colors, including black, brown, red, purple, and green. Each cowpea production region has preferred varieties, valuing certain color and pattern traits above others for determining quality and use. In West Africa, consumers pay a premium for seeds exhibiting certain characteristics specific to the locality, such as lack of color for use as flour or solid brown for use as whole beans [3], [8]–[10]. In the United States consumers prefer varieties with tight black eyes, commonly referred to as “black-eyed peas” [11].

Seed coat traits in cowpea have been studied since the early 20th century, when Spillman [12] and Harland [13], reviewed by Fery [14], explored the inheritance of factors controlling seed coat color and pattern. In a series of F2 populations, Spillman [12] and Harland [13] identified genetic factors responsible for color expression, including “Red” (*R*), which was confirmed by Saunders [15] and Drabo et al [16]. The previous studies noted above, as well as similar studies in soybean (*Glycine max*) [17] and common bean (*Phaseolus vulgaris*) [18]–[20], have shown that there are two loci involved in red coat color, each following a simple Mendelian inheritance. One of those loci seems closely associated with the Color Factor locus, which controls overall pigmentation distribution on the seed coat [19]–[21]. This simple inheritance makes it relatively easy to develop reliable markers for use in marker-assisted breeding. Due to the high level of synteny between cowpea and other leguminous grain crops [22], markers developed in cowpea may be directly useful for breeding in those closely related species.

A genotyping array for 51,128 single nucleotide polymorphisms (SNP) been developed for cowpea [23] which, combined with relatively large populations, offers opportunities to improve the precision of genetic mapping. In particular, new populations have been developed for higher-resolution mapping including a minicore collection representing a worldwide cross-section of cultivated cowpea (“UCR Minicore”) [24] and an eight-parent Multi-parent Advanced Generation Inter-Cross (MAGIC) population [25]. In addition, a reference genome sequence of cowpea (phytozome-next.jgi.doe.gov) [22] and genome assemblies of six additional diverse accessions [26]; (https://phytozome-next.jgi.doe.gov/cowpeapan/) have been produced recently. Here, we make use of genetic and genomic resources available for cowpea to map the red seed coat trait, determine candidate genes, and propose sources of the observed variation.

## 2. Materials and Methods

### 2.1. Plant Materials

Three cowpea populations were used for mapping: the UCR Minicore, which includes 368 accessions [24], a MAGIC population consisting of 305 lines [25], and an F2 population developed as part of this work from a cross between Sasaque (Red, *r-1*) and Sanzi (Not Red, *R-1*), consisting of 108 individuals. Example images of the seed coats, including these two accessions, can be found in **Figure 1**.

**Figure 1.**
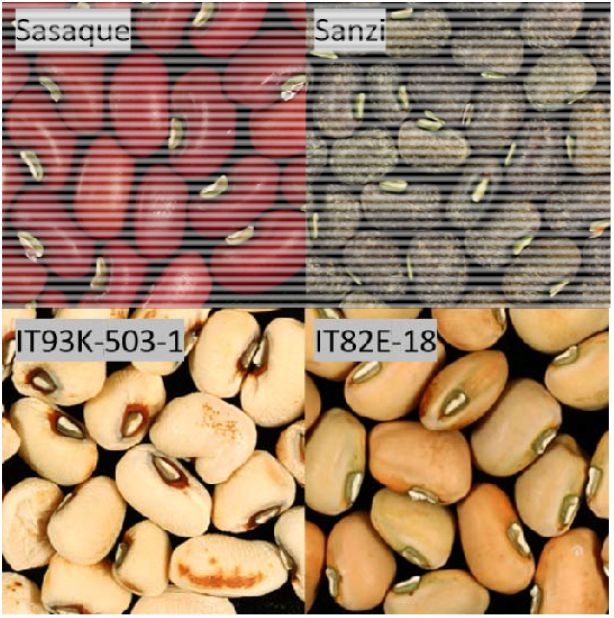
Images of accessions expressing and not expressing red seed coat color. Sasaque and IT93K-503-1 express red seed coat color; Sanzi and IT82E-18 do not.

Candidate gene sequences were compared using the sequences of the cowpea reference genome (IT97K-499-35-1) and the six additional sequences from the pangenome (Sanzi, CB5, Suvita-2, UCR779, ZN016, and TZ30) [26].

These accessions used as parents in the mapping populations and as part of the cowpea pangenome [26] consist of at least one from each of the six identified subpopulations in global cowpea [24] (**Supplementary Table S1**).

### 2.2. SNP Genotyping and Data Curation

DNA was extracted from young leaf tissue using the Qiagen DNeasy Plant Mini Kit (Qiagen, Germany) per the manufacturer’s instructions. The Cowpea iSelect Consortium Array (Illumina Inc., California, USA), which assays 51,128 SNPs [23] was used to genotype each DNA sample. Genotyping was performed at the University of Southern California Molecular Genomics Core facility (Los Angeles, California, USA). The same custom cluster file as in Muñoz-Amatriaín et al [23] was used for SNP calling.

For the UCR Minicore, SNPs with >20% missing data and a minor allele frequency (MAF) < 0.05 were eliminated, leaving a total of 42,603 SNPs used for mapping. For the MAGIC population, SNP data and a genetic map were available from Huynh et al [25], including 32,130 SNPs in 1,568 genetic bins. For the Sasaque by Sanzi F2 population, SNPs were filtered to remove non-polymorphic loci between the parents, leaving 14,772 total polymorphic markers (**Supplementary Table S2**).

### 2.3. Seed Coat Phenotyping

Phenotype data for seed coat traits were collected by visual examination of the seeds. The scored phenotypic classes consisted of “Red” and “Not Red,” as dictated by the presence or absence of observable red pigmentation in the mature seed coat. For mapping purposes, the “Red” lines were marked with a “1” and the “Not Red” with a “0.” Accessions with black seed coat pigmentation or no pigmentation whatsoever were regarded as missing data as it is believed that black pigmentation obscures other colors from view [27] and that the lack of color is controlled by the *Color Factor* locus Herniter et al [21]. Phenotype data can be found in **Supplementary Tables S3** (UCR Minicore), **S4** (MAGIC), and **S5** (Sasaque by Sanzi).

Expected segregation ratios were determined based on the type of population and the parental and, when available, F1 phenotypes. Expected segregation ratios were tested by chi-square analysis.

Dominance relationships were determined by examining the phenotypes of the F1 of the Sasaque by Sanzi population, the segregation of the parental phenotypes in the F2 generation, and individuals from the early development of the MAGIC population. Seeds from relevant individuals were visually examined for the presence of red pigmentation on the seed coat.

### 2.4. Genetic Mapping of the Red Seed Coat Trait

Genetic mapping of the red seed coat trait was achieved with different methods for each type of population. In the MAGIC population, the R package “mpMap” [28] was used as described by Huynh et al [25]. The significance cutoff values were determined through 1000 permutations, resulting in a threshold of *p* = 8.10E-05 [-log10(*p*) = 4.09]. Due to the high number of markers in the genotype data, imputed markers spaced at 1 cM intervals were used.

A Genome-Wide Association Study (GWAS) was performed in the UCR Minicore to identify SNPs associated with the Red phenotype. The weighted mixed-linear model (MLM) function [29] implemented in TASSEL v.5 (www.maizegenetics.net/tassel) was used, with five principal components accounting for population structure in the dataset. The -log10(*p*) values were plotted against the physical coordinates of the SNPs [26]. A Bonferroni correction was applied to correct for multiple testing error in GWAS, with the significance cut-off set at α/n, where α is 0.05 and n is the number of tested markers, resulting in a cut-off of -log10(*p*) = 5.93.

For the Sasaque by Sanzi F2 population, the genotype calls of each bulked DNA pool in the population were filtered to leave only the markers known to be polymorphic between the parents, and these were then sorted based on physical positions in the pseudochromosomes of IT97K-499-35 [26]. The genotype data was then examined visually in Microsoft Excel for areas where the recessive bulk was homozygous and the dominant bulk was heterozygous.

### 2.5. Candidate Gene Identification and PCR Amplification

The gene-annotated sequences of the overlapping QTL and region of interest for both Vu03 and Vu07 were obtained from the v1.2 of the reference genome sequence of cowpea (https://phytozome-next.jgi.doe.gov/info/Vunguiculata_v1_2) [22]. Candidate genes were identified through location within the significant regions and similarity to genes known to be involved in red pigmentation in soybean and common bean in the literature (see Discussion). Additionally, data from the Gene Expression Atlas (VuGEA) of the reference genome was utilized (https://legumeinfo.org) [30].

Primers were developed to amplify sections of the candidate gene *Vigun03g118700* in Sasaque to examine the sequence for causative variations. Each primer pair, as well as the annealing temperatures, are listed in **Supplementary Table S7**. PCR was performed using the Thermo Scientific DreamTaq Green PCR Master Mix (Thermo Scientific, Massachusetts, USA) per the manufacturer’s instructions. Primers were developed using Primer3 v0.4.1 (bioinfo.ut.ee/primer3) and ordered from Integrated DNA Technology (Coralville, Iowa, USA). PCR was run for 25-45 cycles with an annealing temperature compatible with the primer pair and an extension time of 60-75 sec. Amplicons were confirmed by gel electrophoresis. PCR products of the expected size were purified using the QIAquick PCR Purification kit (Qiagen, Germany) and Sanger sequenced in both directions on an Applied Biosystems 3730xl DNA Sequencer (University of California, Riverside IIGB Genomics Core).

Additional sequences of *Vigun03g118700* and *Vigun07g118600* were obtained from IT97K-499-35 (https://phytozome.jgi.doe.gov) and the other genomes sequenced as part of the pangenome, consisting of Sanzi (Red), CB5-2 (Not Red), Suvita-2 (Not Red), UCR779 (Not Red), TZ30 (Red), and ZN016 (Red) (https://phytozome-next.jgi.doe.gov/cowpeapan/) [26]. Complete nucleotide and amino acid sequences of both candidate genes were compared using MultAlin [31].

## 3. Results

### 3.1. Phenotypic Variation for Red Seed Coat Color

As described in Materials and Methods, phenotypic data were collected from the three populations and their parents (when applicable) by visual examination of the seeds. A summary of the phenotypic data, along with predicted segregation ratios, chi-square values, and probability can be found in **Table 1**, with demonstrations in **Figure 2**.

**Table 1.** Seed coat phenotypes and segregation ratios of the tested populations. “Other” includes seeds with black pigmentation or no pigmentation.

| Population (# of lines) | Not Red | Red | Other | Pred. Seg. Ratio | X<sup>2</sup> | Probability |
| --- | --- | --- | --- | --- | --- | --- |
| UCR Minicore (368) | 190 | 80 | 98 | -- | -- | -- |
| MAGIC (305) | 157 | 92 | 56 | 5:3 | 0.032 | 0.86 |
| Sasaque by Sanzi (108) | 86 | 22 | 0 | 3:1 | 1.23 | 0.27 |

**Figure 2.**
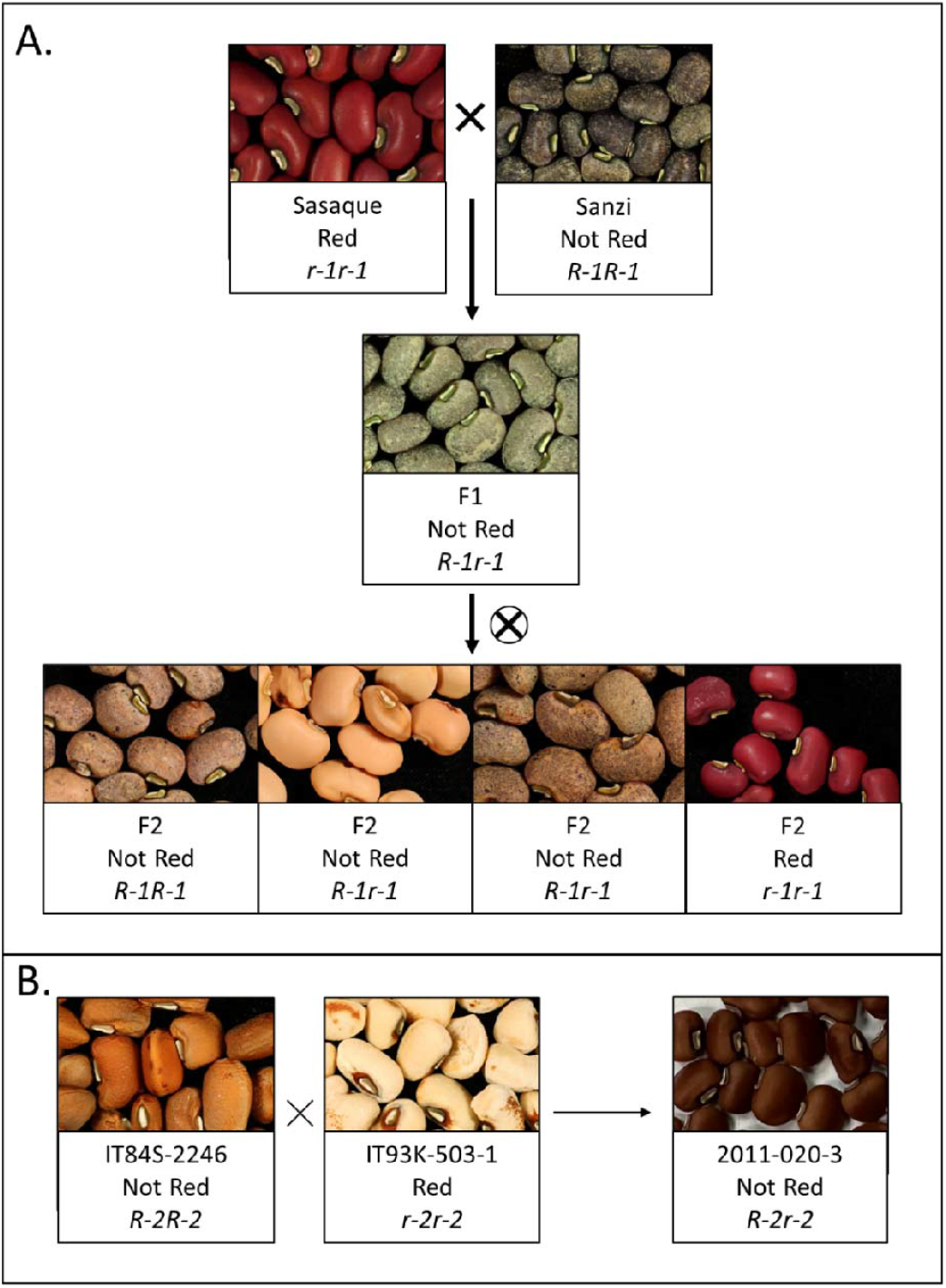
Segregation at the two *Red* loci, *R-1* and *R-2*. (A) Segregation at the *R-1* locus in the Sasaque by Sanzi F2 population. (B) Cross between IT84S-2246 and IT93K-503-1 from the early development of the MAGIC population, resulting in Not Red phenotype in the seed coats on seeds of the F1 maternal parent.

Expected segregation ratios were determined based on the type of population and the parental and, when available, F1 phenotypes. For example, the Sasaque by Sanzi F2 population was expected to segregate in a 3:1 ratio. For the MAGIC population, based on how the population was constructed [25] it was assumed that each fully homozygous parent had a roughly 1/8 chance to pass its genotype at a particular locus to a given RIL. Three of the MAGIC parents had red seed coat (IT89KD-288, IT84S-2049, and IT93K-503-1) while the other five parents did not, resulting in an expected 5:3 segregation ratio.

The observed 3:1 segregation ratio of the Sasaque by Sanzi population together with the observed Not Red seed coat of the F1 indicate the dominance of *R-1* (Not Red) over *r-1* (Red). Similarly, a cross from the early development of the MAGIC population demonstrated the dominance of *R-2* (Not Red) over *r-2* (Red): IT84S-2246 (*R-2R-2*, Not Red) was crossed with IT93K-503-1 (*r-2r-2*, Red). The resulting plant, 2011-020-3 (*R-2r-2*) produced seeds with the Not Red phenotype (**Figure 2B**).

### 3.2. Loci Controlling Red Seed Coat Color

Two QTL, one each on Vu03 and Vu07, were identified using the three populations. The Vu03 QTL was identified in the UCR Minicore and the Sasaque x Sanzi F2 population, while the Vu07 QTL was identified in the UCR Minicore and the MAGIC population (**Table 2; Figure 3**). Higher-resolution mapping for both genomic regions was achieved using the UCR Minicore due to the higher genetic diversity and number of recombination events available in this population. In particular, the significant region was 135 kbp in size for Vu03, and it was reduced to just one SNP position for Vu07 (**Table 2; Supplementary Table S6**). An analysis of linkage disequilibrium (*r*^2^) between pairs of SNPs located 50 kbp upstream and downstream from the significant SNP for Vu07 (2_02375) revealed low LD between the marker and nearby variants (analysis not shown). Complete information on the mapping results can be found in **Supplementary Tables S7** (UCR Minicore) and **S8** (MAGIC).

**Table 2.** Mapping results in the different populations.

| Population | Chr | Peak SNP | Flanking SNPs | QTL pos. (bp) | LOD/-log10 (p) | R<sup>2</sup> (%) | Effect |
| --- | --- | --- | --- | --- | --- | --- | --- |
| Sasaque x Sanzi (F<sub>2</sub>) | Vu03 | -- | 2_38494 - 2_05272 | 7,657,076 - 41,457,212 | -- | -- |  |
| UCR Minicore | Vu03 | 2_20787 | 2_03389 - 2_20787 | 10,933,603 - 11,068,119 | 9.82 | 16.6 | -0.56 (G) |
| UCR Minicore | Vu07 | 2_02375 | -- | 22,014,760 | 6.26 | 9.8 | -0.61 (A) |
| MAGIC | Vu07 | -- | 2_51274 - 2_04829 | 14,976,909 - 22,928,810 | 29.81 | 38.82 |  |

**Figure 3.**
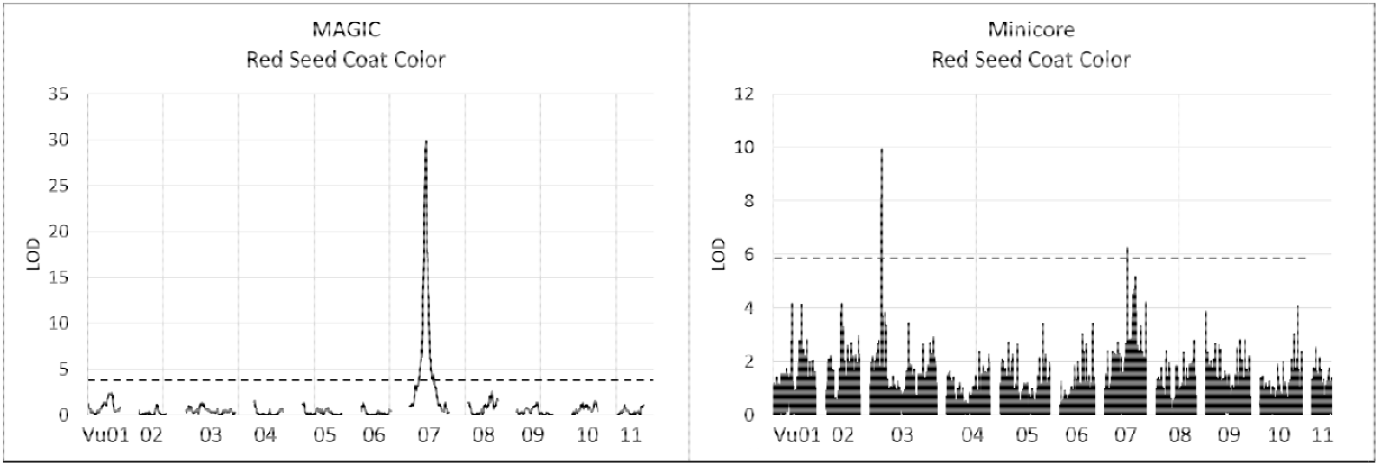
Mapping of the red seed coat trait in the MAGIC and UCR Minicore populations. The significance threshold is indicated by the dashed line.

### 3.3 Identification of Candidate Genes

The overlapping areas of the QTL and regions of interest on Vu03 and Vu07 were examined for candidate genes in the cowpea reference genome v1.2 (phytozome-next.jgi.doe.gov/info/Vunguiculata_v1_2) [22]. Eleven genes were identified in the minimal area (the area in which all the independent mappings identified) on Vu03 and one gene in the minimal area on Vu07 (**Supplementary Table S9**). Of the genes in the minimal area on Vu03, *Vigun03g118700* is the best candidate as it encodes a dihydroflavonol 4-reductase, which shows a highly elevated expression levels during early seed development (**Supplementary Figure S2** [30]). The peak SNP on Vu03 is also located within *Vigun03g118700*. The closest homologs to *Vigun03g118700* are anthocyanidin reductases identified in various leguminous plants, including *Vigna radiata, Vigna angularis*, common bean (*Phaseolus vulgaris*), and soybean (*Glycine max*), among others (**Supplementary Table 10**).

In the UCR Minicore, GWAS only identified a single significant SNP on Vu07, 2_02375, contained within the larger region identified in mapping the MAGIC population (**Table 2, Supplementary Tables S7** and **S8**). This SNP is located in the third intron of *Vigun07g118600*, which is orthologous to the Arabidopsis *BAH1/NLA* gene encoding an E3 ubiquitin-protein ligase with roles in defense response, plant growth and development [32]. The significant SNP is not in high LD with nearby variants. *Vigun07g118600* does not show a high expression in developing seed tissue of the reference genome cv. IT97K-499-35, which possesses black-eyed seeds (https://legumeinfo.org) [30].

To identify possible causative polymorphisms, sequence comparisons were made using the published cowpea genomes [26] and the sequence of *Vigun03g118700* amplified from the Sasaque genome. Variant locations in the following data are calculated from the start of the coding sequence.

In *Vigun03g118700*, several SNPs were observed among accessions, one of which could be causative (**Figure 4, Table 3**). In the two Red-seeded genotypes from the pangenome, TZ30 and ZN016, there is a substitution at base 593, C*→*A, resulting in an amino acid substitution, Ala*→*Asp. In Sasaque, there were two substitutions. One at base 661, T*→*G, resulting in an amino acid substitution, Met*→*Ser; the other at base 672, G*→*T, resulting in an amino acid substitution, Val*→*Phe.

**Table 3.** Observed variations in the coding DNA sequence of the candidate genes. The base number is calculated from the start of the coding sequence. “849+” indicates that including base 849 to the end of the coding sequence.

| Vigun03g118700 |  |  |  |  |  |  |
| --- | --- | --- | --- | --- | --- | --- |
| <u>BP</u> | <u>RefGen Base</u> | <u>Mutant Base</u> | <u>Genotypes Present In</u> | <u>AA Code</u> | <u>AA</u> | <u>Mutation Type</u> |
| 378 | C | T | UCR779 | ATT→ATC | Ile→Ile | Silent |
| 592 | G | A | UCR779 | ACT→GCT | Thr→Ala | Polar Uncharged → hydrophobic |
| 593 | C | A | ZN016, TZ30 | GCT→GAT | Ala→Asp | hydrophobic → polar negative |
| 661 | T | G | Sasaque | ATG→AGC | Met→Ser | hydrophobic → polar uncharged |
| 672 | G | T | Sasaque | GTT→TTT | Val→Phe | hydrophobic → hydrophobic |
| Vigun07g118600 |  |  |  |  |  |  |
| <u>BP</u> | <u>RefGen Base</u> | <u>Mutant Base</u> | <u>Genotypes Present In</u> | <u>AA Code</u> | <u>AA</u> | <u>Mutation Type</u> |
| 761 | T | C | UCR779 | TTC→TCC | Phe→Ser | hydrophobic → polar uncharged |
| 849+ |  |  | ZN016, TZ30 |  |  | Completely different AA sequence |

**Figure 4.**
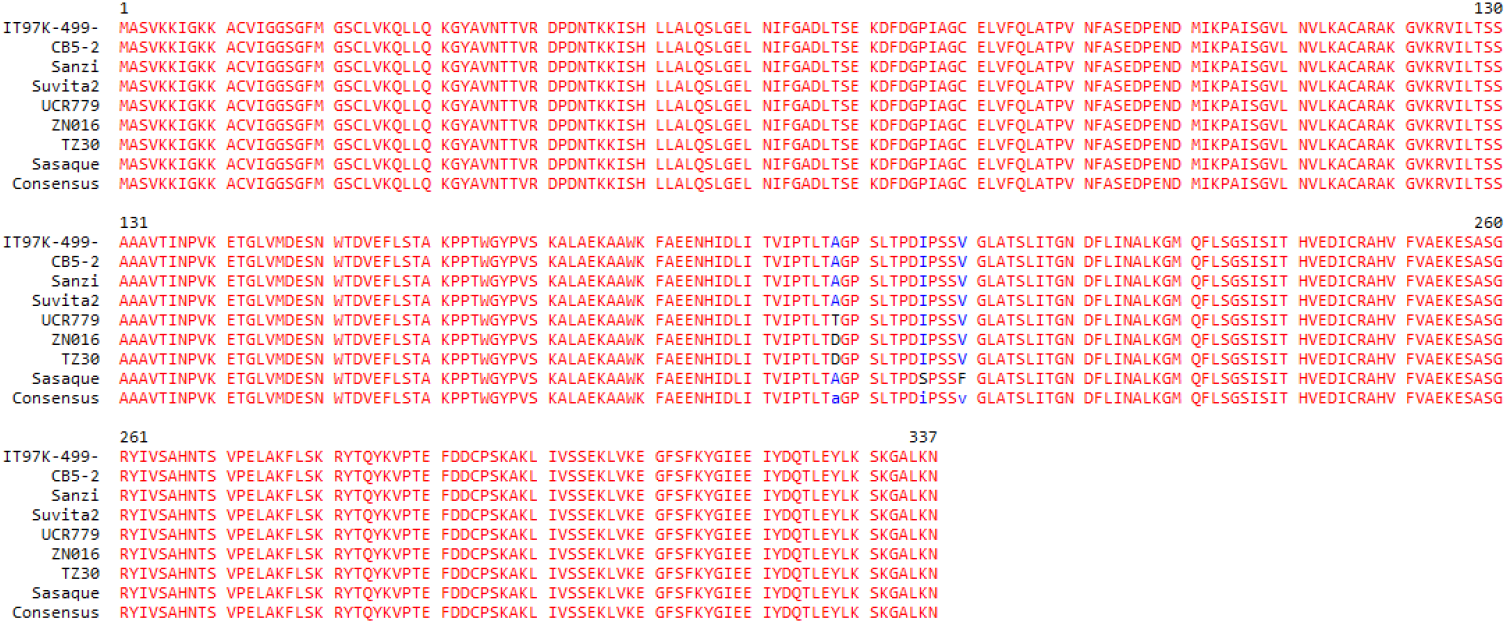
Alignment of proposed amino acid sequences of *Vigun03g118700* among genotypes in the pangenome with the addition of Sasaque. Sasaque, ZN016, and TZ30 express the red seed coat phenotype. The other genotypes do not have red seed coat coloring.

In addition, there were two substitutions in UCR779, a brown-seeded genotype from the pangenome. At base 378, there is a silent mutation, C*→*T, still encoding Ile; and at base 592, G*→*A, resulting in an amino acid substitution, Thr-Ala.

In *Vigun07g118600*, additional mutations were observed (Figure 5, Table 3). In both TZ30 and ZN016 there are major differences from base 850 onwards, resulting in a different amino acid sequence. In addition, at base 761 in UCR779, there is a SNP, T*→*C, resulting in an amino acid substitution.

**Figure 5.**
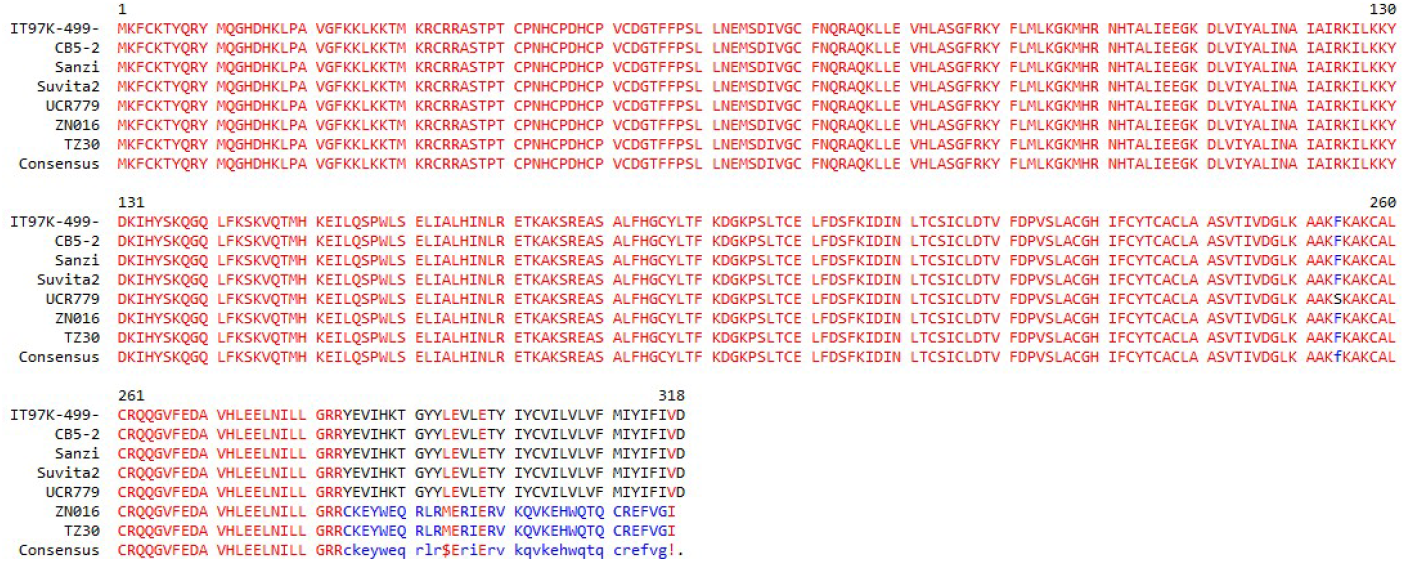
Alignment of proposed amino acid sequences of *Vigun07g118600* among genotypes in the pangenome. ZN016 and TZ30 express the red seed coat phenotype. The other genotypes do not have red seed coat coloring.

## 4. Discussion

Previous studies, dating back over a hundred years, by Harland [13] and Spillman [12], reviewed by Fery [14] and confirmed by Drabo et al [16], identified only the single locus *R* as being responsible for the red seed coat phenotype in cowpea. The present data suggests the presence of two loci each of which can be independently responsible for the expression of red seed coat color. To avoid confusion, the locus on Vu03 is referred to as *R-1* while the locus on Vu07 is referred to as *R-2*. While the previous studies in cowpea and soybean have only identified a single locus responsible for red seed coat color, Bassett et al [33] identified two loci in common bean, termed *Rk* (red kidney) and R (oxblood red). The candidate gene on Vu07, *Vigun07g118600*, is closely linked the Color Factor locus, believed to be controlled by *Vigun07g110700* [21], just 1.5 Mb [26] and 0.96 cM distant according to the consensus map established by Muñoz-Amatriaín et al [23]. This occurrence could explain why a second locus for Red color in cowpea was not previously identified, as on seeds with only small areas of pigmentation, such as IT93K-503-1 (**Figure 1**) it can be difficult to discern Red (*r-1*/*r-1, r-2*/*r-2*) from Not Red (*R-1, R-2*). A similar linkage has been well-documented in common bean (*Phaseolus vulgaris*) [33], and has been recently confirmed through trait mapping [19], [20].

The flavonoid biosynthesis pathway has been well characterized in a number of plants, including soybean [17] and Arabidopsis [34], among others. An intermediate group of molecules in the flavonoid biosynthesis pathway are cyanidins. Cyanidins are well-described pigmentation molecules, identified in a wide range of plants, including red cabbage [35], clover [36], and various red berries [37], among others. Cyanidins appear red in low pH (<3.5) conditions [38]. It has been well-established much of the observed variation in plant organ pigmentation arises from differential levels of pigmentation molecules and pH in those organs [39].

The present study reports the identification of the *R-1* locus on Vu03. The most likely candidate gene, *Vigun03g118700*, encodes a dihydroflavonol 4-reductase protein (D4R). Based on the dominance pattern, with Not Red (*R-1*) dominant towards Red (*r-1*), it is most likely that this is the previously identified *R* locus from the literature [14], [16]. A BLAST search of the soybean (*Glycine max*) genome on Phytozome, showed that the closest homolog of *Vigun03g118700* in soybean *Glyma.08G062000* (previously referred to as *Glyma08g06630*), with 90.63% identity and a e-value of 0.0 from pairwise BLAST of the coding sequence.

Kovinich et al [17] demonstrated that soybean mutants with reduced function of the soybean *ANR1* gene (*Glyma.08G062000*) had a red seed coat phenotype. This would be consistent with a complete dominance model as a single functional copy of the gene would allow successful catalysis of cyanidin to epicatechin and result in no red pigmentation, while an individual with two recessive alleles and nonfunctional D4R would not catalyze the biosynthesis of epicatechin, resulting in a buildup of cyanidin, and instead would express red seed coat pigmentation. The observed allele sequence in common bean would match this, with the dominant *Rk* allele not expressing red pigmentation, the recessive *rk*^*d*^ allele expressing red pigmentation, and the semidominant *rk* allele in the middle, with some red pigmentation [18].

Similarly, the closest homolog of *Vigun03g118700* in common bean is *Phvul.002G218700*, with 95.66% identity and an e-value of 0.0 from pairwise BLAST of the coding sequences. This gene is also predicted to encode an ANR. The recent mapping of color in common bean by Campa et al [19] and Sadohara et al [20] identified SNPs which were significantly associated with the *a\** trait, which measures greenness-redness, as determined by the International Commission on Illumination [40]. Both found significantly correlated markers located 6-15 Mb up and downstream of *Phvul.002218700*. It is possible that the mapping did not identify markers closer because they did not map Red versus Not Red phenotypes directly, incorporating the green color as part of the phenotyping.

While the sequence comparisons of *Vigun03g118700* identified several potentially causative variations in Sasaque, TZ30, and ZN016, they also identified variations present in UCR779, which does not have any red pigmentation (Table 3). While the base substitution at base 378 (C*→*T, Ile*→*Ile) in *Vigun03g118700* is a silent mutation, the substitutions at base 592 (G*→*A, Thr*→*Ala) in *Vigun03118700* and at base 761 (T*→*C, Phe*→*Ser) in *Vigun07g118600* are more substantial. It is unclear why these mutations do not result in a similarly expected loss of the function to the mutations found in the Red phenotyped varieties. Perhaps some insight could be gained from the determination that UCR779 is part of the most highly divergent subpopulation of cowpea [24], from eastern and southern Africa. It may be that its sequence represents alternative Not Red alleles (*r-1 & R-2*). To better understand this, future studies should examine other diverse cowpea genotypes.

The *R-2* locus identified in the present study mapped to Vu07. Like the dominant red R locus in common bean, Not Red color was dominant at this locus (**Figure 2B**) and it is closely linked to the Color Factor (C) locus, previously mapped by Herniter et al [21]. The lone significant SNP identified here falls within an intron of *Vigun07g118600*, which is predicted to encode an E3 ubiquitin ligase. The available expression data for this gene shows quite low expression, with only about 0.5 TPM in the early developing seed and dropping off thereafter (**Supplementary Figure S3**). However, it should be noted that the only genotype for which transcription data is available is the cowpea reference genome, IT97K-499-35, which does not have red pigmentation and likely has the recessive, non-functional (*r-2*) allele, which is likely not expressed. This expression differential might be explained by variation in the promoter region. Future studies should seek to identify the transcription factor which controls expression of *Vigun07g118600* and identify potential binding sites which could vary between genotypes.

While the function of *Vigun03g118700* can be inferred based on previous research in other species, the function of *Vigun07g118600* in seed coat color is much less clear. Some possibilities include regulation of the flavonoid biosynthesis pathway, perhaps even exerting negative control over D4R (*R-1* locus, *Vigun03g118700*). Similarly, the closest homolog in soybean is *Glyma.10G018800*, with 90.26% identity and an e-value of 0.0 from pairwise BLAST of the coding sequence. Interestingly, *Glyma.10G018800* shows elevated expression in the root (https://phytozome-next.jgi.doe.gov/report/transcript/Gmax_Wm82_a4_v1/Glyma.10G018800.1) and has been identified as a candidate gene controlling root growth [41], [42], potentially resulting from divergent evolution.

The closest homolog of *Vigun07g118600* in common bean is *Phvul.007G143600*. In their mappings, both Sadohara et al [20] and Campa et al [19] identified a SNP 6 Mb downstream of *Phvul.007G143600*, as significantly associated with the *a\** trait. However, Sadohara et al [20] did not proffer candidate genes and Campa et al [19] did not consider this gene. It may be that, similarly to the function of the homolog in soybean, the function of *Phvul.007G143600* has diverged over evolutionary time.

## 5. Conclusions

This study presents the mapping of red seed coat color in cowpea, identifying two loci, one each on Vu03 and Vu07, and proposing a candidate gene for each, *Vigun03g118700*, encoding a dihydroflavonol 4-reductase, and *Vigun07g118600*, encoding an E3 ubiquitin ligase. Sequence comparisons between accessions either expressing or not expressing red pigmentation identified multiple potentially causative variations. The red seed coat color is likely caused by the disruption of the flavonoid biosynthesis pathway, resulting in a buildup of anthocyanidin in the seed coat.

## Supplementary Materials

The following are available online at www.mdpi.com/xxx/s1, Figure S1: *Vigun03g118700* expression profile, Figure S2: *Vigun07g118600* expression profile, Figure S3: Coding sequence alignment of *Vigun03g118700* among the pangenome sequences, Figure S4: Coding sequence alignment of *Vigun07g118600* among the pangenome sequences, Table S1: Country of origin and subpopulation affinity of the population parents and pangenome accessions, Table S2: Genotypes of the bulked samples from the F2 population Sasaque by Sanzi, Table S3: Phenotypes of the UCR Minicore Collection, Table S4: Phenotypes of the UCR MAGIC population, Table S5: Phenotypes of the F2 Sasaque by Sanzi population, Table S6: Primers used to amplify the candidate gene *Vigun03g118700* for sequencing, Table S7: GWAS results from the UCR Minicore Collection, Table S8: Mapping results from the UCR MAGIC population, Table S9: Candidate genes identified within the QTL peaks, Table S10: BLASTP Results of the *Vigun03g118700* transcript.

## Author Contributions

Conceptualization, I.A.H., T.J.C.; data curation, I.A.H., St.L.; formal analysis, I.A.H., M.M.A.; funding acquisition, St.L., T.J.C.; investigation, I.A.H., M.M.A., Sa.L., and Y.N.G.; methodology, I.A.H.; project administration, I.A.H., M.M.A., and T.J.C.; resources, I.A.H., T.J.C.; software, St.L.; supervision, T.J.C.; validation, I.A.H., M.M.A.; visualization, I.A.H., M.M.A.; writing – original draft, I.A.H.; writing – review and editing, I.A.H., St.L., M.M.A., Sa.L., Y.N.G., and T.J.C. All authors have read and agreed to the published version of the manuscript.

## Informed Consent Statement

Not applicable.

## Data Availability Statement

The original contributions presented in the study are included in the article/supplementary material; further inquiries can be directed to the corresponding author.

## Acknowledgments

The authors thank Eric Castillo, Mia Rochford, Julia Valdepeña, and Sabrina Phengsy for assistance with seed photography; Steve Wanamaker for assistance in the analysis of the various genome sequences; Bao-Lam Huynh for providing the MAGIC population and attendant genotypic and pedigree information; Pei Xu and Xinyi Yu for providing images of the TZ30 and ZN016 genotypes.

## References

[1] FAOSTAT, “Crops,” 2023. http://www.fao.org/faostat/en/#data/QC (accessed Feb. 26, 2020).

[2] J. D. Ehlers and A. E. Hall, “Cowpea (Vigna unguiculata L. Walp.),” F. Crop. Res., vol. 53, no. 1–3, pp. 187–204, 1997, doi: 10.1016/S0378-4290(97)00031-2.

[3] O. Boukar et al., “Cowpea (Vigna unguiculata): Genetics, genomics and breeding,” Plant Breed., vol. 138, no. 4, pp. 415–424, Aug. 2019, doi: 10.1111/pbr.12589.

[4] B. B. Singh, Cowpea: The Food Legume of the 21st Century. 2014. Accessed: May 11, 2016. [Online]. Available: https://dl.sciencesocieties.org/publications/books/tocs/acsesspublicati/cowpeathefoodle

[5] A. R. Tijjani, R. T. Nabinta, and M. Muntaka, “Adoption of innovative cowpea production practices in a rural area of Katsina State, Nigeria,” J. Agric. Crop Res., vol. 3, no. June, pp. 53–58, 2015.

[6] S. R. Jaeger, L. Antúnez, G. Ares, M. Swaney-Stueve, D. Jin, and F. R. Harker, “Quality perceptions regarding external appearance of apples: Insights from experts and consumers in four countries,” Postharvest Biol. Technol., vol. 146, pp. 99–107, Dec. 2018, doi: 10.1016/J.POSTHARVBIO.2018.08.014.

[7] A. S. Kostyla, F. M. Clydesdale, and M. R. McDaniel, “The psychophysical relationships between color and flavor,” Food Sci. Nutr., vol. 10, no. 3, pp. 303–321, 1978, doi: 10.1080/10408397809527253.

[8] A. S. Langyintuo et al., “Cowpea supply and demand in West and Central Africa,” F. Crop. Res., vol. 82, no. 2, pp. 215–231, 2003, doi: 10.1016/S0378-4290(03)00039-X.

[9] F. J. Mishili et al., “Consumer preferences for quality characteristics along the cowpea value chain in Nigeria, Ghana, and Mali,” Agribusiness, vol. 25, no. 1, pp. 16–35, Sep. 2009, doi: 10.1002/agr.20184.

[10] A. H. Ira, J. Zhenyu, and K. Francis, “Market preferences for cowpea (Vigna unguiculata [L.] Walp) dry grain in Ghana,” African J. Agric. Res., vol. 14, no. 22, pp. 928–934, May 2019, doi: 10.5897/AJAR2019.13997.

[11] R. L. Fery, “The Genetics of Cowpeas: A Review of the World Literature,” in Cowpea Research, Production and Utilization, S. R. Singh and K. O. Rachie, Eds. New York: John Wiley & Sons, Inc., 1985, pp. 25–62.

[12] W. J. Spillman, “Inheritance of the ‘eye’ in Vigna,” Am. Nat., vol. XLV, no. 53, pp. 513–523, 1911.

[13] S. C. Harland, “Inheritance of certain characters in the cowpea (Vigna sinensis),” J. Genet., vol. 8, no. 2, pp. 101–132, Apr. 1919, doi: 10.1007/BF02983490.

[14] R. L. Fery, “Genetics of Vigna,” in Horticultural Reviews, vol. 2, Hoboken, NJ, USA: John Wiley & Sons, Inc., 1980, pp. 311–394. doi: 10.1002/9781118060759.ch7.

[15] A. R. Saunders, “Inheritance in the cowpea (Vigna sinensis Endb.). II: seed coat colour pattern; flower, plant, and pod color,” South African J. Agric. Sci., vol. 3, no. 2, pp. 141–162, 1960.

[16] I. Drabo, T. A. O. Ladeinde, J. B. Smithson, and R. Redden, “Inheritance of Eye Pattern and Seed Coat Colour in Cowpea (Vigna unguiculata [L.] Walp.),” Plant Breed., vol. 100, no. 2, pp. 119–123, Mar. 1988, doi: 10.1111/j.1439-0523.1988.tb00226.x.

[17] N. Kovinich, A. Saleem, J. T. Arnason, and B. Miki, “Identification of two anthocyanidin reductase genes and three red-brown soybean accessions with reduced anthocyanidin reductase 1 mRNA, activity, and seed coat proanthocyanidin amounts,” J. Agric. Food Chem., vol. 60, no. 2, pp. 574–584, Jan. 2012, doi: 10.1021/jf2033939.

[18] M. J. Bassett, “Genetics of Seed Coat Color and Pattern in Common Bean,” in Plant Breeding Reviews, 28th ed., Hoboken, NJ, USA: Wiley, 2007, pp. 239–315. doi: 10.1002/9780470168028.ch8.

[19] A. Campa, R. Rodríguez Madrera, M. Jurado, C. García-Fernández, B. Suárez Valles, and J. J. Ferreira, “Genome-wide association study for the extractable phenolic profile and coat color of common bean seeds (Phaseolus vulgaris L.),” BMC Plant Biol., vol. 23, no. 1, pp. 1–16, Dec. 2023, doi: 10.1186/S12870-023-04177-Z/TABLES/5.

[20] R. Sadohara, Y. Long, P. Izquierdo, C. A. Urrea, D. Morris, and K. Cichy, “Seed coat color genetics and genotype × environment effects in yellow beans via machine-learning and genome-wide association,” Plant Genome, vol. 15, no. 1, p. e20173, Mar. 2021, doi: 10.1002/tpg2.20173.

[21] I. A. Herniter et al., “Seed Coat Pattern QTL and Development in Cowpea (Vigna unguiculata [L.] Walp.),” Front. Plant Sci., vol. 10, p. 514455, Oct. 2019, doi: 10.3389/fpls.2019.01346.

[22] S. Lonardi et al., “The genome of cowpea (Vigna unguiculata [L.] Walp.),” Plant J., vol. 98, no. 5, pp. 767–782, Jun. 2019, doi: 10.1111/tpj.14349.

[23] M. Muñoz-Amatriaín et al., “Genome resources for climate-resilient cowpea, an essential crop for food security,” Plant J., vol. 89, no. 5, pp. 1042–1054, Mar. 2017, doi: 10.1111/tpj.13404.

[24] M. Muñoz-Amatriaín et al., “The UCR Minicore: a valuable resource for cowpea research and breeding,” Legum. Sci., p. leg3.95, May 2021, doi: 10.1002/leg3.95.

[25] B.-L. Huynh et al., “A multi-parent advanced generation inter-cross (MAGIC) population for genetic analysis and improvement of cowpea (Vigna unguiculata L. Walp.),” Plant J., vol. 93, no. 6, pp. 1129–1142, Mar. 2018, doi: 10.1111/tpj.13827.

[26] Q. Liang et al., “A view of the pan-genome of domesticated Cowpea (Vigna unguiculata [L.] Walp.),” Plant Genome, p. e20319, Mar. 2023, doi: 10.1002/TPG2.20319.

[27] I. A. Herniter, M. Muñoz-Amatriaín, S. Lo, Y.-N. Guo, and T. J. Close, “Identification of Candidate Genes Controlling Black Seed Coat and Pod Tip Color in Cowpea (Vigna unguiculata [L.] Walp),” G3; Genes|Genomes|Genetics, vol. 8, no. 10, pp. 3347–3355, Oct. 2018, doi: 10.1534/g3.118.200521.

[28] B. E. Huang and A. W. George, “R/mpMap: a computational platform for the genetic analysis of multiparent recombinant inbred lines,” Bioinformatics, vol. 27, no. 5, pp. 727–729, Mar. 2011, doi: 10.1093/bioinformatics/btq719.

[29] Z. Zhang et al., “Mixed linear model approach adapted for genome-wide association studies,” Nat. Genet., vol. 42, no. 4, pp. 355–360, Apr. 2010, doi: 10.1038/ng.546.

[30] S. Yao et al., “The Vigna unguiculata Gene Expression Atlas (VuGEA) from de novo assembly and quantification of RNA-seq data provides insights into seed maturation mechanisms,” Plant J., vol. 88, no. 2, pp. 318–327, Oct. 2016, doi: 10.1111/tpj.13279.

[31] F. Corpet, “Multiple sequence alignment with hierarchical clustering,” Nucleic Acids Res., vol. 16, no. 22, pp. 10881–10890, Nov. 1988, doi: 10.1093/NAR/16.22.10881.

[32] T. Yaeno and K. Iba, “BAH1/NLA, a RING-type ubiquitin E3 ligase, regulates the accumulation of salicylic acid and immune responses to Pseudomonas syringae DC3000,” Plant Physiol., vol. 148, no. 2, pp. 1032–1041, 2008, doi: 10.1104/pp.108.124529.

[33] M. J. Bassett, R. Lee, C. Otto, and P. E. Mcclean, “Classical and Molecular Genetic Studies of the Strong Greenish Yellow Seedcoat Color in ‘Wagenaar’ and ‘Enola’ Common Bean,” J. Am. Soc. Hortic. Sci., vol. 127, no. 1, pp. 50–55, 2002.

[34] K. Petroni and C. Tonelli, “Recent advances on the regulation of anthocyanin synthesis in reproductive organs,” Plant Sci., vol. 181, no. 3, pp. 219–229, 2011, doi: 10.1016/j.plantsci.2011.05.009.

[35] N. Ahmadiani, R. J. Robbins, T. M. Collins, and M. M. Giusti, “Molar absorptivity (ε) and spectral characteristics of cyanidin-based anthocyanins from red cabbage,” Food Chem., vol. 197, pp. 900–906, Apr. 2016, doi: 10.1016/j.foodchem.2015.11.032.

[36] H. Zhang, H. Tian, M. Chen, J. Xiong, H. Cai, and Y. Liu, “Transcriptome analysis reveals potential genes involved in flower pigmentation in a red-flowered mutant of white clover (Trifolium repens L.),” Genomics, vol. 110, no. 3, pp. 191–200, May 2018, doi: 10.1016/j.ygeno.2017.09.011.

[37] M. J. Jara-Palacios, A. Santisteban, B. Gordillo, D. Hernanz, F. J. Heredia, and M. L. Escudero-Gilete, “Comparative study of red berry pomaces (blueberry, red raspberry, red currant and blackberry) as source of antioxidants and pigments,” Eur. Food Res. Technol., vol. 245, no. 1, pp. 1–9, Jan. 2019, doi: 10.1007/s00217-018-3135-z.

[38] S. Asen, K. H. Norris, and R. N. Stewart, “Absorption spectra and color of aluminium-cyanidin 3-glucoside complexes as influenced by pH,” Phytochemistry, vol. 8, no. 3, pp. 653–659, Mar. 1969, doi: 10.1016/S0031-9422(00)85416-3.

[39] J. Mol, E. Grofewold, and R. Koes, “How genes paint flowers and seeds,” Trends in Plant Science, vol. 3, no. 6. pp. 212–217, Jun. 01, 1998. doi: 10.1016/S1360-1385(98)01242-4.

[40] CIE, “Commission Internationale de l’Eclairage CIE 015: 2018 Colorimetry,” 2018. doi: 10.25039/TR.015.2018.

[41] S. Sarkar, A. Shekoofa, A. McClure, and J. D. Gillman, “Phenotyping and Quantitative Trait Locus Analysis for the Limited Transpiration Trait in an Upper-Mid South Soybean Recombinant Inbred Line Population (‘Jackson’ × ‘KS4895’): High Throughput Aquaporin Inhibitor Screening,” Front. Plant Sci., vol. 12, p. 779834, Jan. 2022, doi:10.3389/FPLS.2021.779834/BIBTEX.

[42] X. Li, X. Zhang, Q. Zhao, and H. Liao, “Genetic improvement of legume roots for adaption to acid soils,” Crop J., vol. 11, no. 4, pp. 1022–1033, Aug. 2023, doi: 10.1016/J.CJ.2023.04.002.

